# Dkk1 inhibition normalizes limb phenotypes in a mouse model of *Fzd2* associated omodysplasia Robinow syndromes

**DOI:** 10.1101/2022.06.02.494481

**Authors:** Ryan P. Liegel, Megan N. Michalski, Sanika Vaidya, Elizabeth Bittermann, Erin Finnerty, Chelsea A. Menke, Cassandra R. Diegel, Zhendong A. Zhong, Bart O. Williams, Rolf W. Stottmann

## Abstract

FRIZZLED-2 (FZD2) is a transmembrane Wnt ligand receptor. We previously identified a pathogenic human *FZD2* variant, encoding for a protein with a premature stop and loss of 17 amino acids. This includes a portion of the consensus DISHEVELLED binding sequence required for Wnt signal transduction. Patients with this variant exhibited FZD2-associated autosomal dominant Robinow Syndrome. To model this variant, we utilized zygote microinjection and *i*-GONAD-based CRISPR/Cas9-mediated genome editing to generate an allelic series in the mouse. Embryos mosaic for humanized *Fzd2*^*W553\**^ knock-in exhibited cleft palate and shortened limbs, consistent with *FZD2*^*W548\**^ patient phenotypes. We also generated two germline mouse alleles with small deletions, *Fzd2*^*D3*^ and *Fzd2*^*D4*^. Homozygotes for each allele survive embryonic development at normal ratios but exhibit a highly penetrant cleft palate phenotype, shortened limbs compared to wild-type and perinatal lethality. *Fzd2*^*D4*^ craniofacial tissues indicated decreased canonical WNT signaling. *In utero* treatment with IIIC3a (DKK inhibitor) normalized the limb lengths in *Fzd2*^*D4*^ homozygotes. The *in vivo* replication represents an approach to further investigate the mechanism of FZD2 phenotypes and validates the utility of CRISPR knock-in mice as a tool for demonstrating pathogenicity of human genetic variants. We also present evidence for a potential therapeutic intervention.

## INTRODUCTION

*FRIZZLED-2* encodes a transmembrane receptor which is an essential component in Wnt signal transduction (1-5). FZD2 is part of a diverse family of 10 FRIZZLED receptors that interact with 19 WNT ligands. Wnt signaling regulates a variety of developmental processes throughout all tissues in the body through a combination of spatio-temporal regulation of signaling molecule expression and differential ligand-receptor interactions. Many human syndromes are caused by the disruption of *WNT* or *FZD*, including autosomal dominant omodysplasia (ADO) (6-11) and autosomal dominant Robinow syndrome (AD-RS) (12-17); yet the mechanisms underlying these syndromes need further investigation in model organisms.

Our previous findings demonstrated the first evidence of *FZD2* variants in human pathology (8). In two related individuals, a single nucleotide variant leading to a premature stop allele in *FZD2* (W548*) resulted in ADO, a syndrome characterized by small stature due to the shortening of long bones, craniofacial dysmorphism including cleft lip/palate, and genitourinary abnormalities. Subsequently, other reports have supported the conclusion that heterozygous variants in *FZD2* can result in limb and craniofacial anomalies. Turkmen et al. reported a missense variant, G434V, in a patient with ADO (10). Shortly after, Nagasaki et al. identified a patient with a premature stop variant at the amino acid position directly upstream of W548, S547* (7). Additional W548* and G434V variants were reported by White et al. (17) and Warren et al. (11) in patients with AD-RS and ADO. White et al. (17) also identified G434S and W377* variants, and Zhang et al. reported another three related individuals with the W548* variant as well as a F130Cfs*98 variant and G434S (18).

Due to the highly overlapping phenotypes seen in patients classified with ADO and AD-RS, there has been some confusion as to how patients with FZD2-related skeletal phenotypes should be classified. Zhang et al. performed a Human Phenotype Ontology analysis for 16 subjects with FZD2 variants and concluded all FZD2 patients grouped with patients that had known variants in other genes associated with Robinow syndrome (18). They suggested that patients with Fzd2 variants that result in characteristic limb shortening and craniofacial malformations be classified as FZD2-associated AD-RS. We agree with this suggestion and use this nomenclature moving forward.

While our report (8) was the first *FZD2* human variant to be identified, two previous studies had investigated FZD2 loss-of-function in a mouse model (19, 20). They reported a partially penetrant phenotype of recessive cleft palate (50%) as well as incompletely penetrant failure to thrive in the animals without cleft palate. While these studies are consistent with the human FZD2-associated AD-RS phenotype, there are differences. The *Fzd2* knockout animals were reported to have hypognathia but were not reported to have limb reductions. Also, the homozygous knockout animals display more mild craniofacial phenotypes than the ROR2 knockout animals which have been reported to phenocopy Robinow Syndrome (21). We hypothesized these differences might be due to the comparison of a *Fzd2* null mouse model with a dominant human syndrome. These studies were unable to definitively determine if the effects on Wnt signaling primarily compromised the canonical and/or non-canonical portions of the pathway. We therefore sought to further investigate the role of the *FZD2* W548* patient variant in mice to identify if disruption of only the C-terminal region of *Fzd2* in mouse was phenotypically different than a complete null. The C-terminal region of FZD2 is highly conserved between mouse and human which makes modeling the pathogenic allele in mouse particularly relevant to human FZD2 biology. Here, we present the generation of the mouse ortholog of this patient allele of FZD2 as well as two additional lines containing small deletions in the same C-terminal region of the protein.

We present data that the orthologous W553* allele in mice recapitulates the autosomal dominant inheritance pattern seen in human patients and that both the skeletal and craniofacial abnormalities from the human patients were observed in mice. We also present data in which the canonical and noncanonical WNT pathways are perturbed in these mouse mutants. Augmenting the canonical WNT signaling pathway resulted in some rescue of several skeletal phenotypes.

## RESULTS

### Development of Fzd2 C-terminal tail variant mouse models

To model the human *FZD2*^*W548\**^ variant, we utilized CRISPR-Cas9-mediated genome editing to create the orthologous W553* change in mice (*Fzd2*^*W553\**^). C57BL6/J zygotes were injected with sgRNAs and a DNA oligonucleotide donor. Sanger sequencing of the targeted region in mosaic founder animals revealed the presence of editing for the desired W553* knock-in as well as multiple indels (**Figure 1A-B**).

**Figure 1.**
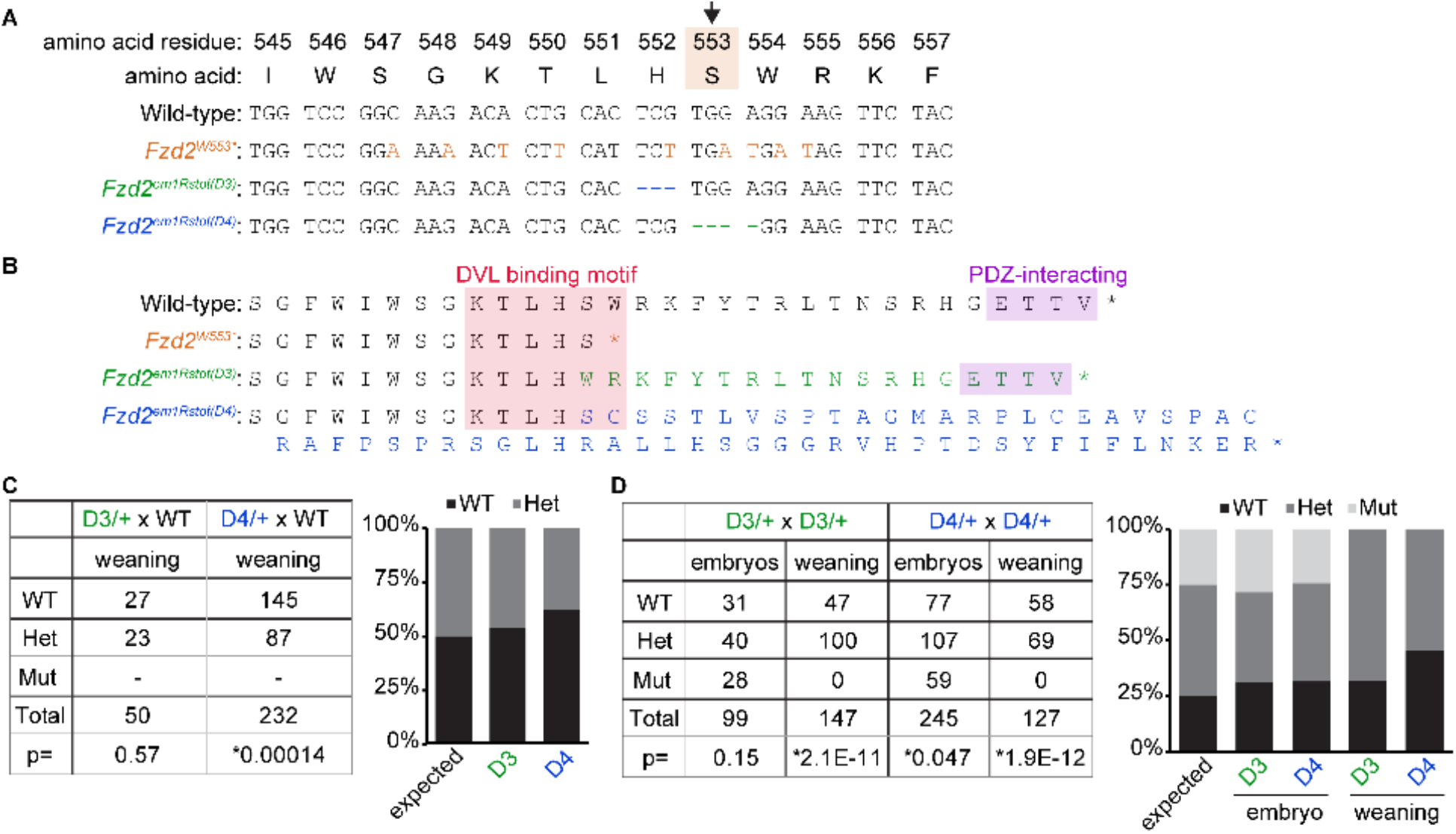
Mouse models with FZD2 C-terminal modifications cause perinatal lethality. (**A** and **B**) Zygote microinjection of CRISPR/CAS9 reagents targeting the C-terminal region to generate the orthologous human *FZD2*^*W548\**^ variant in mice (*Fzd2*^*W553\**^) resulted in three different mouse models: the target variant *Fzd2*^*W553\**^ and two unintended indels *Fzd2*^*em1Rstot(D3)*^ and *Fzd2*^*em1Rstot(D4)*^. (**C**) Quantification of WT and heterozygous weanlings from *Fzd2*^*D3/+*^ or *Fzd2*^*D4/+*^ crossed with WT animals. (**D**) Quantification of WT, heterozygous, and homozygous embryos (E17.5) or P28 weanlings from heterozygous by heterozygous crosses for both the *Fzd2*^*D3/+*^ and *Fzd2*^*D4/+*^ lines.

Crossing the mosaic founders to wild-type C57BL6/J mice resulted in the generation of stable lines of two small deletions.

*Fzd2*^*em1Rstot(D3)*^ is a 3bp deletion precisely removing the codon for Serine 552 (S552) and is hereafter referred to as *Fzd2*^*D3*^ (**Figure 1A-B**). *Fzd2*^*em2Rstot(D4)*^ is a 4bp deletion that results in an immediate frameshift at amino acid 552 and a stop codon following 60 residues of nonsense sequence and is hereafter referred to as *Fzd2*^*D4*^ (**Figure 1A-B**). While there was some evidence of low levels of editing for W553* in founders, we were unable to recover this specific allele in any offspring to generate a stable line. All the alleles we generated are predicted to disrupt a portion of the canonical Dishevelled binding domain (KTxxxW, **Figure 1B**) (22).

In maintenance of the *Fzd2*^*D3*^ and *Fzd2*^*D4*^ alleles by mating *Fzd2* carriers with wild-type (WT) mice, we noted normal proportions of *Fzd2*^*D3/+*^ mice but a significant reduction in *Fzd2*^*D4/+*^ mice at weaning (**Figure 1C**). Heterozygous matings of both the *Fzd2*^*D3/+*^ and *Fzd2*^*D4/+*^ alleles revealed that both *Fzd2*^*D3/D3*^ and *Fzd2*^*D4/D4*^ homozygous embryos were recovered in approximately expected Mendelian ratios at late embryonic time points (E16.5-18.5) (**Figure 1D**). We saw a slight reduction in the number of *Fzd2*^*D4/D4*^ homozygote embryos (**Figure 1 D**); however, no *Fzd2*^*D3/D3*^ or *Fzd2*^*D4/D4*^ homozygous animals survived to weaning (**Figure 1D**).

### Heterozygous W553* variants lead to perinatal lethality due to fully penetrant cleft palate

We were not able to create a stable mouse line that recapitulates the orthologous human variant, *Fzd2*^*W553*/+*^. This was expected as heterozygotes for the W553* variant were predicted to have cleft palate, which leads to perinatal lethality in mice because of an inability to feed. We therefore performed an additional round of CRISPR-Cas9 zygotic injections in addition to several rounds of *i*mproved Genome Editing of Oviductal Nucleic Acid Delivery (*i-*GONAD) (23, 24) to generate additional animals for phenotypic analyses. In these “F0” experiments, we collected the resulting embryos at E16.5-18.5 to investigate phenotypes in animals that otherwise would be unavailable at weaning age due to the cleft palate phenotype.

From the second round of zygote injections, 20 embryos were recovered at E17.5 and Sanger sequencing contained evidence of genome editing for seven. Of these seven, five were mosaic (50-90%) for the desired W553* heterozygous knock in (**Figure 2A**), and two were mosaic for multiple other indels which did not create the W553* edit.

**Figure 2.**
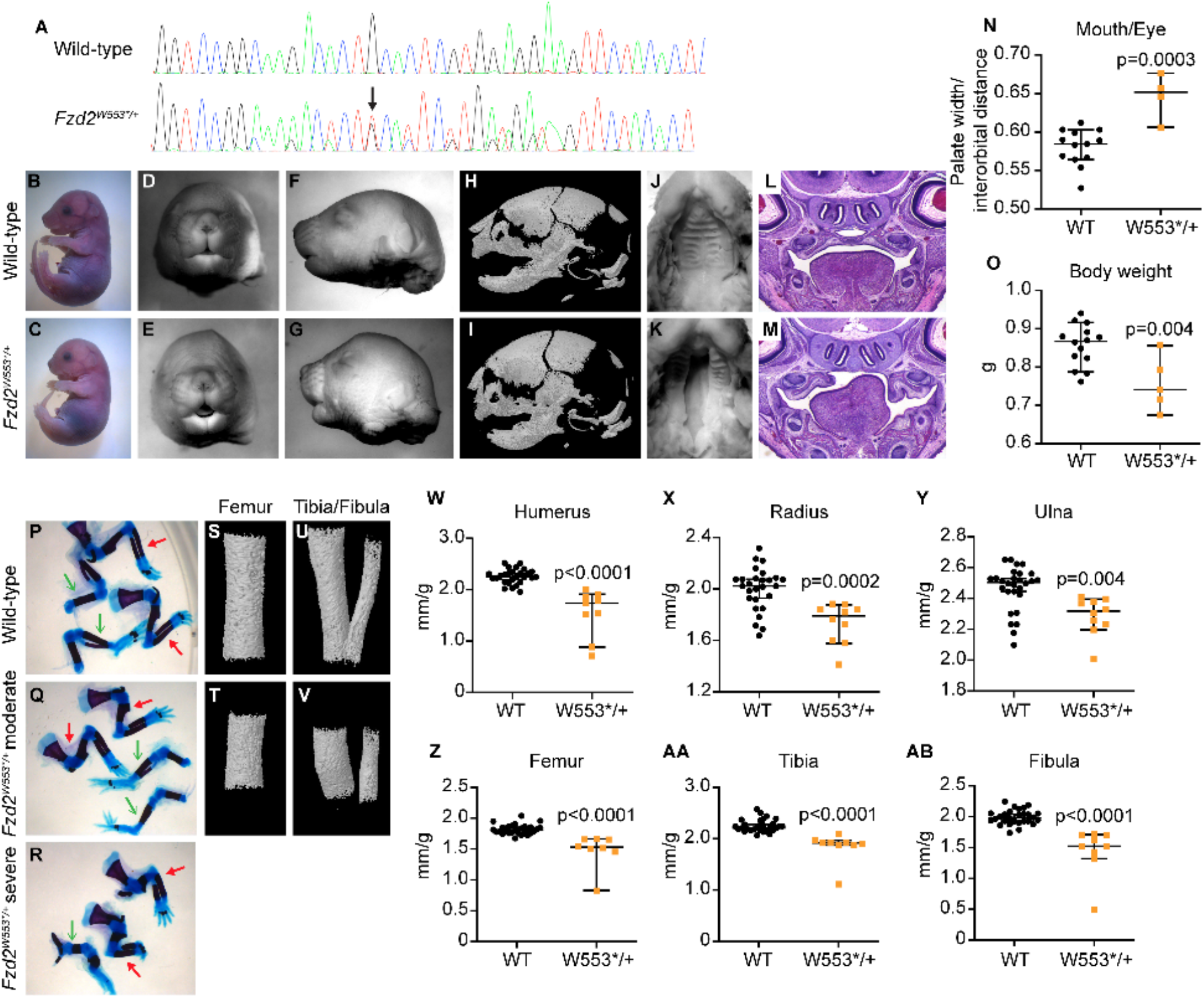
*Fzd2*^*W553*/+*^ embryos have fully penetrant cleft palate and limb shortening. (**A**) Sanger sequencing chromatograms of wild-type and *Fzd2*^*W553*/+*^ edited embryos. (**B-C**) Full-body images of E17.5 wild-type and W553* heterozygous embryos. (**D-G**) Front and left side view images of E15.5 wild-type and W553* heterozygous heads. (**H-I**) 3D Micro-CT images of skulls from E18.5 wild-type and W553* heterozygous heads. (**J-K**) Images of E17.5 wild-type and W553* heterozygous palates. (**L-M**) Coronal sections through the mid-palate of E17.5 wild-type and W553* heterozygous embryos. (**N**) Width of the mouth normalized to the distance between the eyes n=4-13. (**O**) Body weight measurements. n=5-14/group. (**P-R**) Images of forelimb (red closed arrows) and hindlimb (green open arrows) skeletal preps from wild-type and W553* heterozygotes with moderate and severe limb shortening. (**S-V**) 3D micro-CT images of femur (**S**,**T**) or tibia/fibula (**U**,**V**) from wild-type and W553* heterozygotes. (**W-AB**) Length measurements of humeri, radii, ulnae, femora, tibiae, and fibulae (n=2 per animal) per embryo body weight. All p values are indicated on graphs.

### Heterozygous W553* variants recapitulate human FZD2-associated AD-RS phenotypes

All *Fzd2*^*W553*/+*^ -edited embryos had short frontonasal prominences (**Figure 2C, G, I**) relative to wild-type littermate controls (**Figure 2B, F, H**). The mouth opening was wider in *Fzd2*^*W553*/+*^ -edited embryos (**Figure 2E**) as compared to littermates (**Figure 2D, O**). Examination of the palates of these embryos revealed that all five embryos with the W553* variant had cleft palates, whereas none of the unedited embryos or embryos with other indels had cleft palates (**Figure 2J-M**). Thus, we conclude that the presence of the *Fzd2*^*W553*/+*^ allele is sufficient to cause a “dominant” cleft palate phenotype despite the presence of some WT *Fzd2* sequence via Sanger sequencing (**Figure 2A**). The lower face was significantly wider in *Fzd2*^*W553*/+*^ - edited embryos relative to interorbital width (**Figure 2N**).

In addition to facial dysmorphism and cleft palate, *Fzd2*^*W553*/+*^-edited embryos weighed significantly less than their littermate controls (**Figure 2O**). Because limb shortening is the most dramatic phenotype seen in human FZD2-associated AD-RS, we performed skeletal preparation staining on all “F0” embryos to measure limb lengths. This skeletal analysis revealed that the lengths of the humeri, radii, ulnae, femora, tibiae, and fibulae were significantly shorter in *Fzd2*^*W553*/+*^-edited embryos as compared to littermate controls, even by normalizing to embryo weight to control for size differences (**Figure 2P-AB**). While all *Fzd2*^*W553*/+*^-edited embryos had shortened limbs, there was some variability in the severity of the shortening, with some embryos having very severe loss of bone structure (**Figure 2R**). Overall, skeletal staining showed that the *Fzd2*^*W553*/+*^ allele has a dominant effect on bone length.

To efficiently generate additional embryos with the W553* variant to further confirm the penetrance of phenotypes, we utilized *i*-GONAD which allows for the direct injection of CRISPR-CAS9 reagents into the oviduct of a pregnant female mouse followed by oviductal electroporation. Using this technique, we generated an additional 7 animals containing the Fzd2W553* modification and all animals had cleft palate, further confirming complete penetrance of the phenotype. Additionally, several embryos with unintended insertion/deletion alleles were generated and had cleft palate (**Supplemental Table 1**). In frame insertions did not affect phenotypes, however small to large deletions near amino acid residue 553 resulted in embryos with cleft palate. Additionally, several missense variants led to cleft palate.

### Craniofacial malformations in *Fzd2*^*D3/D3*^ and *Fzd2*^*D4/D4*^ mice

To determine the cause of the lethality in *Fzd2*^*D3/D3*^, *Fzd2*^*D4/D4*^ and *Fzd2*^*D4/+*^ animals, we examined embryos throughout embryonic development (**Figure 3A-I, Supplemental Figure 1A-C**). Histological examination of embryos revealed an incompletely penetrant cleft palate in *Fzd2*^*D3/D3*^ embryos (**Supplemental Figure 1C-E**, n=25/31). However, cleft palate was fully penetrant in all *Fzd2*^*D4/D4*^ embryos (**Figure 3I-K**, n=59/59). None of the WT, *Fzd2*^*D4/+*^, or *Fzd2*^*D3/+*^ heterozygous embryos exhibited palate closure or elevation defects (summarized in **Figure 3K, Supplemental Figure 1D**). Measurement of the palate width as normalized to interorbital distance revealed that *Fzd2*^*D4/D4*^ and cleft *Fzd2*^*D3/D3*^ embryos had significantly wider palates than *Fzd2*^*D4/+*^, *Fzd2*^*D3/+*^, *Fzd2*^*D4/+*^ or non-cleft *Fzd2*^*D3/D3*^ embryos (**Figure 3J, Supplemental Figure 1E**). This is grossly visible in some *Fzd2*^*D4/D4*^ embryos at E13.5 (**Figure 3C arrows**). This finding suggests that the widening of the lower face is highly correlated with cleft palate status. As cleft palate results in perinatal lethality in mice, it is unsurprising that there are no surviving *Fzd2*^*D4/D4*^ animals at weaning. However, another mechanism must be responsible for the postnatal lethality of the few non-cleft *Fzd2*^*D3/D3*^ mice as well as the fraction of *Fzd2*^*D4/+*^ animals that do not survive to weaning. In addition to palatal defects, *Fzd2*^*D4/D4*^ had significantly shorter mandibles compared to WT controls (**Figure 3L-O**). Interestingly, *Fzd2*^*D3/D3*^ non-cleft animals had shorter mandibles, but animals with cleft palates had similar length mandibles relative to controls (**Supplemental Figure 1F-I**).

**Figure 3.**
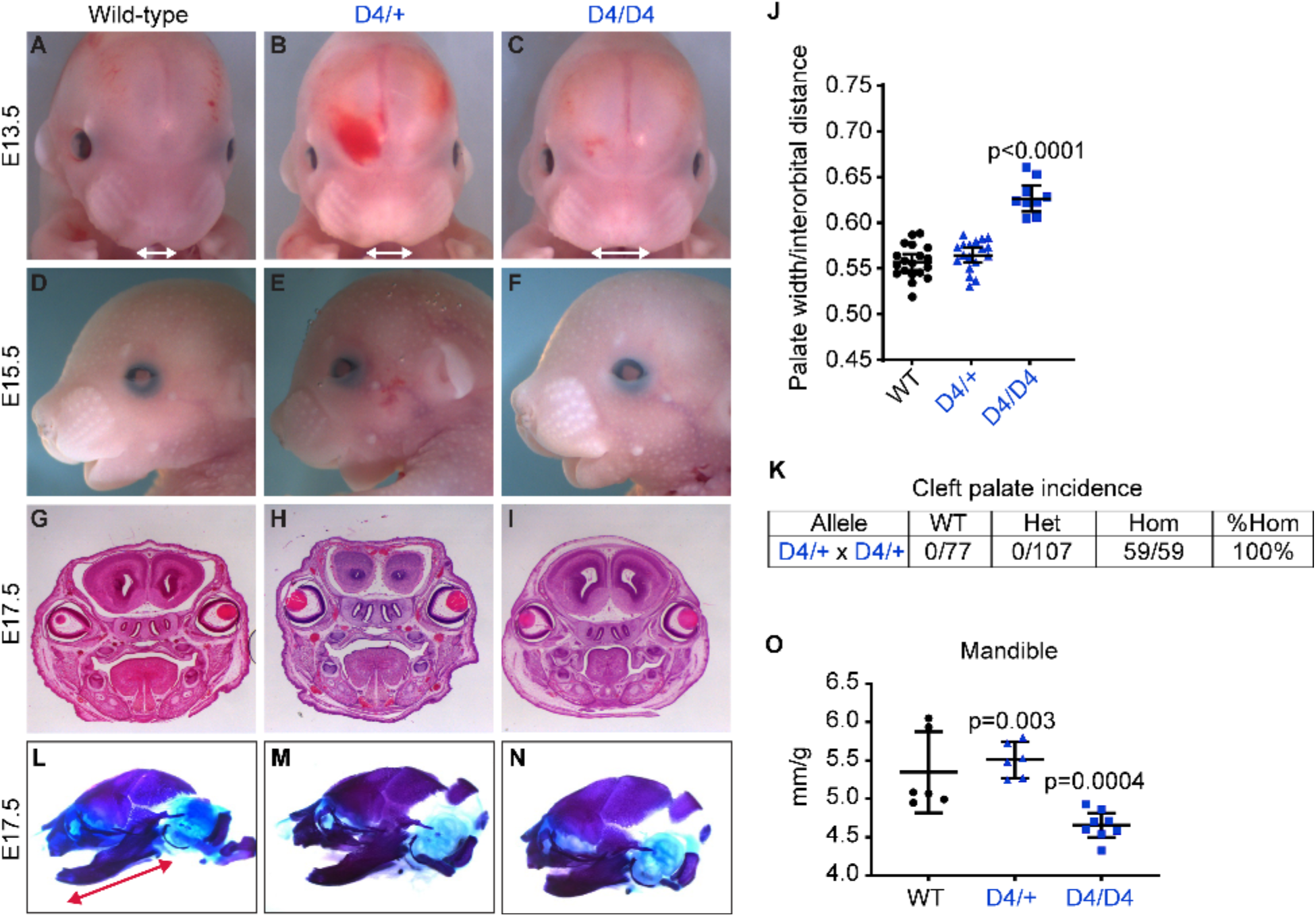
*Fzd2*^*em1Rstot(D4)*^ homozygous animals have increased palatal width and associated fully penetrant cleft palate with shortened mandibles. (**A-C**) Images of E13.5 wild-type, heterozygous and homozygous D4 animals. Arrows highlighting the width of the palate. (**D-F**) Left side view images of E15.5 wild-type, heterozygous and homozygous D4 animals. (**G-I**) Coronal sections through the mid-palate of E17.5 wild-type, heterozygous and homozygous D4 animals. (**J**) Palatal width as a percentage of interorbital width. n=9-21/group. (**K**) Incidence of cleft palate in E17.5 embryos recovered from D4/+ x D4/+ matings. (**L-N**) Left side view images of skeletal preps of skulls from wild-type, heterozygous and homozygous E17.5 D4 animals. Cartilage is stained blue and bone is stained purple. Red double-ended arrow in panel **L** indicates the plane in which the mandible was measured. (**O**) Mandible length (n=2 bones per animal) per body weight. All p values are indicated on graphs.

### Modifications in the C-terminus of FZD2 resulted in shortened limb bones

Patients with FZD2-associated AD-RS have shortened limb elements. To determine whether *Fzd2*^*D3*^ or *Fzd2*^*D4*^ animals had limb phenotypes, E17.5 embryos were stained with Alcian blue (cartilage) and Alizarin red (bone), and the bone lengths were measured as a fraction of body weight. All limb elements from *Fzd2*^*D3/D3*^ and *Fzd2*^*D4/D4*^ embryos were statistically significantly decreased in length (**Figure 4A-J, Supplemental Figure 2A-J)**. *Fzd2*^*D4/+*^ animals had limb lengths consistent with WT animals, whereas *Fzd2*^*D3/+*^ embryos had statistically significant reduction in limb lengths, but not to the degree of *Fzd2*^*D3/D3*^ animals.

**Figure 4.**
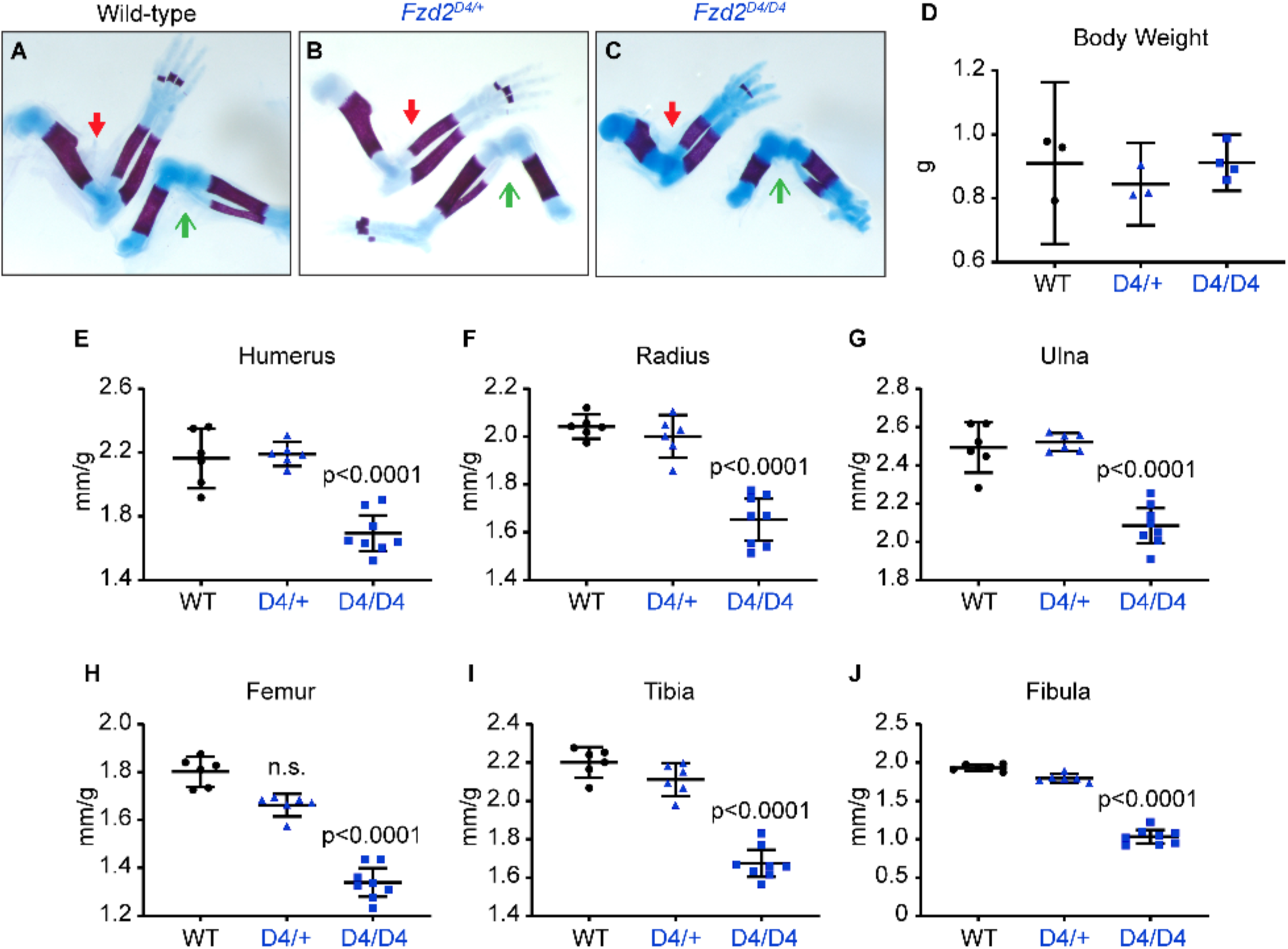
Limb elements are shorter in E17.5 *Fzd2*^*em1Rstot(D4)*^ homozygotes. (**A-C**) Whole mount images of skeletal preps of forelimbs (red closed arrows) and hindlimbs (green open arrows) from E17.5 wild-type, heterozygous and homozygous D4 animals. (**D**) Body weight measurements. n=3-4/group. (**E-J**) Length measurements of humeri, radii, ulnae, femora, tibiae, and fibulae (n=2 per animal) per embryo body weight.

### Genetic background contributions to phenotype

The previously published *Fzd2*^*null*^ mouse was generated on the Sv129 background strain (19) and had a different phenotype than the CRISPR alleles generated in this study on a C57BL/6J (B6) background. We investigated the possibility that genetic background was contributing to the differing phenotypic severity in the two studies. We crossed *Fzd2*^*D4/+*^ mice to wild-type CD-1 mice, an outbred strain with many sequence variants as compared to B6. In this F1 generation, there were increased numbers of *Fzd2*^*D4/+*^ (45%) as opposed to only 37% from the B6 experiment (**Figure 5A, B**), suggesting that even a 50% contribution of the CD-1 outbred strain partially rescues the lethality phenotype seen in a significant proportion of the *Fzd2*^*D4/+*^ animals on the B6 background. Upon mating the heterozygous F1 B6/CD-1 *Fzd2*^*D4/+*^ mice together, *Fzd2*^*D4/D4*^ homozygosity was still lethal after birth. However, embryonic evaluation revealed that 8/10 F1 B6/CD-1 *Fzd2*^*D4/D4*^ animals had cleft palate (**Figure 5C**), as opposed to the 100% penetrant cleft palate on the C57BL6/J background. Interestingly, a small portion (3/26) of the F1 B6/CD-1 *Fzd2*^*D4/+*^ heterozygous embryos developed cleft palate (**Figure 5C**). Together, these data show that the phenotypes resulting from the *Fzd2*^*D4*^ allele with a more dramatic disruption of the C-terminal region of FZD2 remain generally consistent, but with slightly increased variability, on mixed genetic backgrounds. Therefore, we conclude the differences we note in comparison to the null allele are likely due to different molecular mechanisms of action with the C-terminal modifications as compared to complete deletion of *Fzd2*.

**Figure 5.**
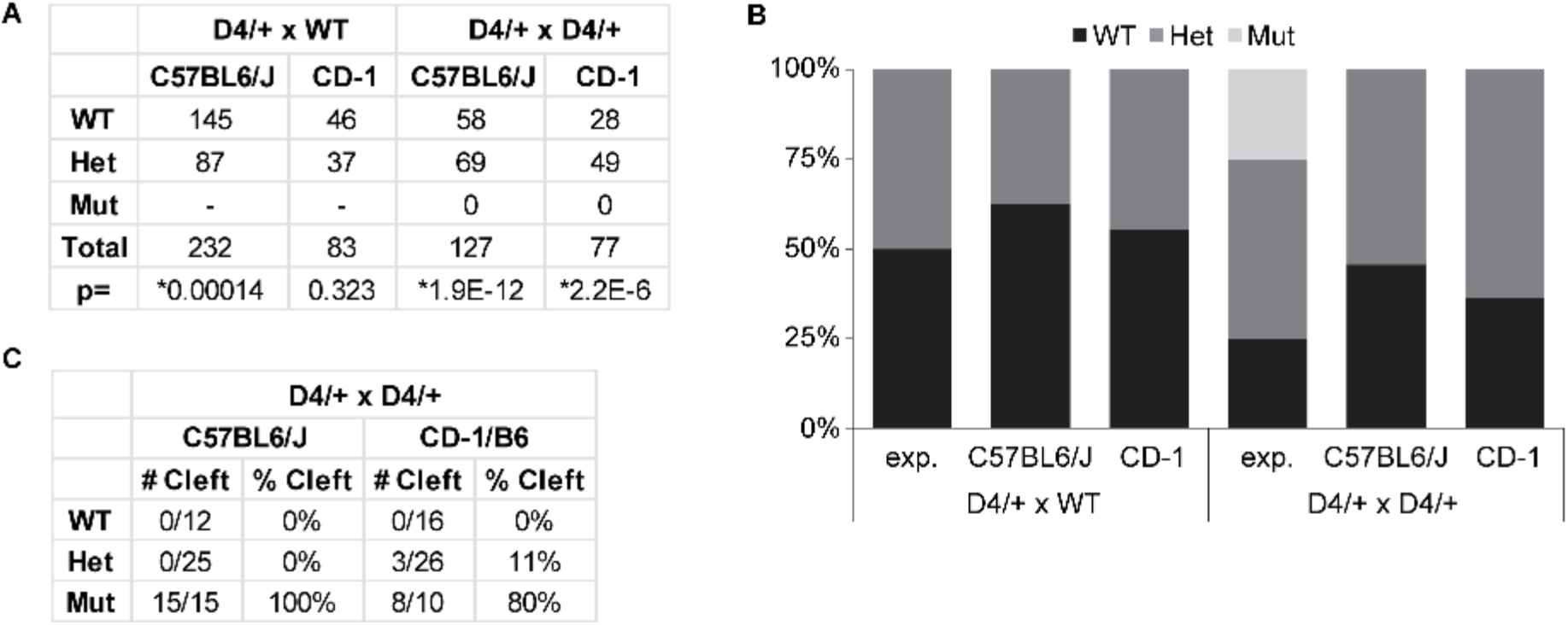
Genetic background affects incidence of cleft palate in *Fzd2*^*em1Rstot(D4)*^ mice. (**A, B**) Quantification of WT and heterozygous weanlings from *Fzd2*^*D4/+*^ crossed with WT animals or WT, heterozygous, and homozygous or weanlings from heterozygous by heterozygous *Fzd2*^*D4*^ crosses on the C57BL6/J or CD-1 background. (**C**) Incidence of cleft palate in E17.5 embryos recovered from D4/+ x D4/+ matings on either the C57BL6/J or C57BL6/J-CD-1 mixed background.

#### Palatal cell populations had normal proliferation and apoptosis

To better understand whether the craniofacial phenotypes observed in the *Fzd2*^*D4/D4*^ animals were related to changes in cell proliferation or cell death, we examined embryonic palatal shelves at E13.5 for proliferation (phospho-histone H3) and apoptotic cell death (cleaved caspase 3).

Immunohistochemistry analysis did not identify a significant difference in either proliferation or apoptosis at E13.5 (**Figure 6A,B**), suggesting that the palatal phenotype is not due to changes in cellular proliferation/death.

**Figure 6.**
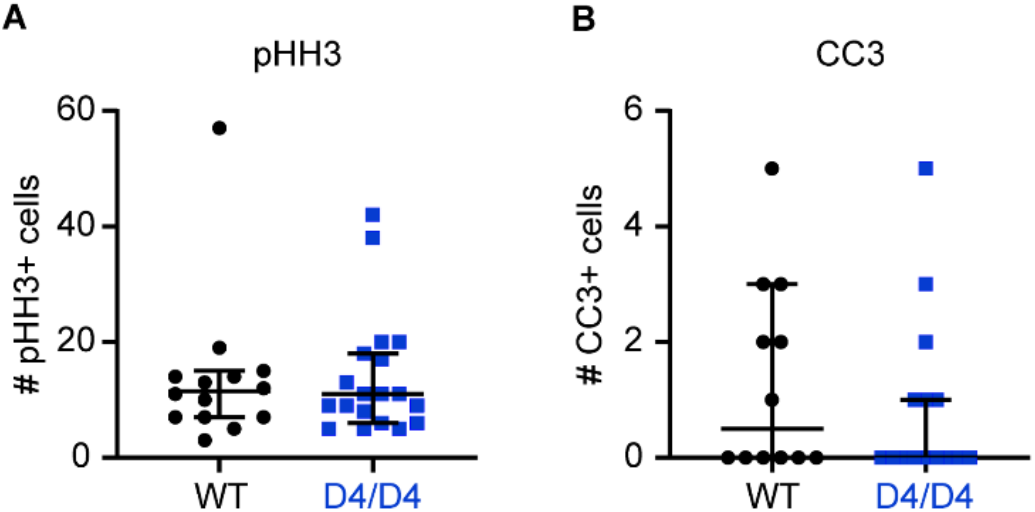
Palatal shelves from *Fzd2*^*em1Rstot(D4)*^ mice did not display differences in cellular proliferation or apoptosis. (**A**) Palatal shelves from E13.5 WT and *Fzd2*^*D4/D4*^ were assessed for changes in cellular proliferation via immunohistochemistry staining for pHH3. The total number of pHH3 positive cells per palatal shelf is plotted. Two to three palatal shelf images were quantified from each WT (n=5) and *Fzd2*^*D4/D4*^ (n=5) animal and are plotted as individual data points. (**B**) Quantification of cellular apoptosis via immunohistochemistry for CC3. Two to three palatal shelf images were quantified from each WT (n=5) and *Fzd2*^*D4/D4*^ (n=5) animal and are plotted as individual data points.

### Alterations in Wnt signaling were observed in tissues from *Fzd2*^*D4/D4*^ embryos

The mouse models generated in this study confirm that disruptions of the cytoplasmic tail of the FZD2 protein are sufficient to recapitulate the FZD2-associated AD-RS phenotypes first identified in human patients (8) and subsequently reported in other patients (7, 10, 11, 17, 18). The ability to maintain the D4 line as heterozygotes allows for generation of additional embryos to explore the underlying molecular mechanisms. The effects on both canonical and non-canonial/PCP Wnt signaling are of particular interest as previous efforts to explore this question have suggested that both pathways may be affected (19, 20).

We first performed qRT-PCR on craniofacial tissues from E13.5 *Fzd2*^*D4/D4*^ embryos for transcriptional targets of canonical Wnt signaling: *Axin2* and *Dkk1*. We found that both are reduced (**Figure 7A, B**) and conclude that the disruptions of the FZD2 cytoplasmic tail prevent full binding of the DVL protein and disrupt normal effects on the β-catenin destruction complex.

**Figure 7.**
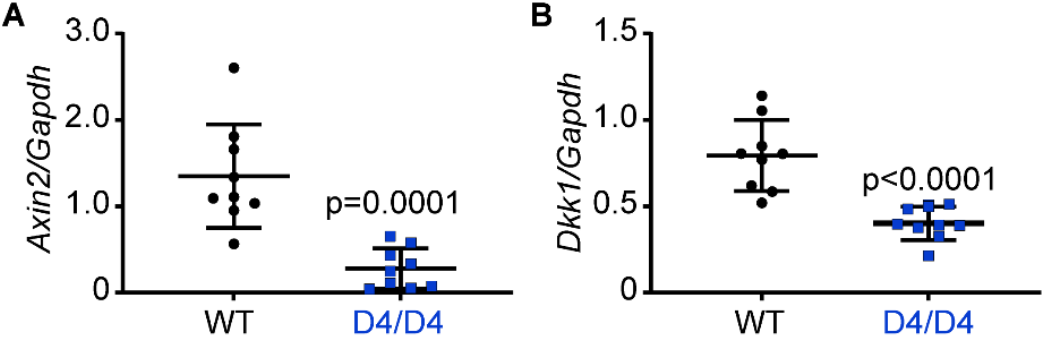
Palatal shelves from *Fzd2*^*em1Rstot(D4)*^ homozygotes display altered canonical WNT signaling. (**A-B**) RNA was isolated from E13.5 wild-type and *Fzd2*^*em1Rstot(D4)*^ homozygotes. *Axin2* (**A**) and *Dkk1* (**B**) mRNA expression levels were quantified relative to GAPDH. n=3 biological replicates/group with 3 technical replicates each. p values are indicated on graphs.

We then addressed the hypothesis that non-canonical Wnt signaling is affected. This is more challenging to assess as there are less well-established parameters to measure this form of Wnt signaling. We measured the length and width of the chondrocytes in the E13.5 limb as a measure of the cell’s ability to position itself within a field of growing cells. Previous reports have linked perturbations of non-canonical Wnt signaling with a reduction in the cells ability to lengthen parallel to the longitudinal axis of limb growth (25-28). We first measured the length and width of chondrocytes with wheat germ agglutinin and found an approximately 50% reduction in the length/width ratio of the cells (**Figure 8A-C**). We repeated the experiment with Safranin-O staining with similar results (**Figure 8D-I**). We conclude from these data that both canonical and non-canonical Wnt signaling are disrupted in FZD2 mutant tissues. Additionally, MEFs generated from *Fzd2*^*W553\**^ embryos showed altered baseline non-canonical signaling. Namely, phospho-VANGL2 was significantly decreased in MEFs from FZD2^W553*^ animals compared to WT controls. (**Figure 8J, K**).

**Figure 8.**
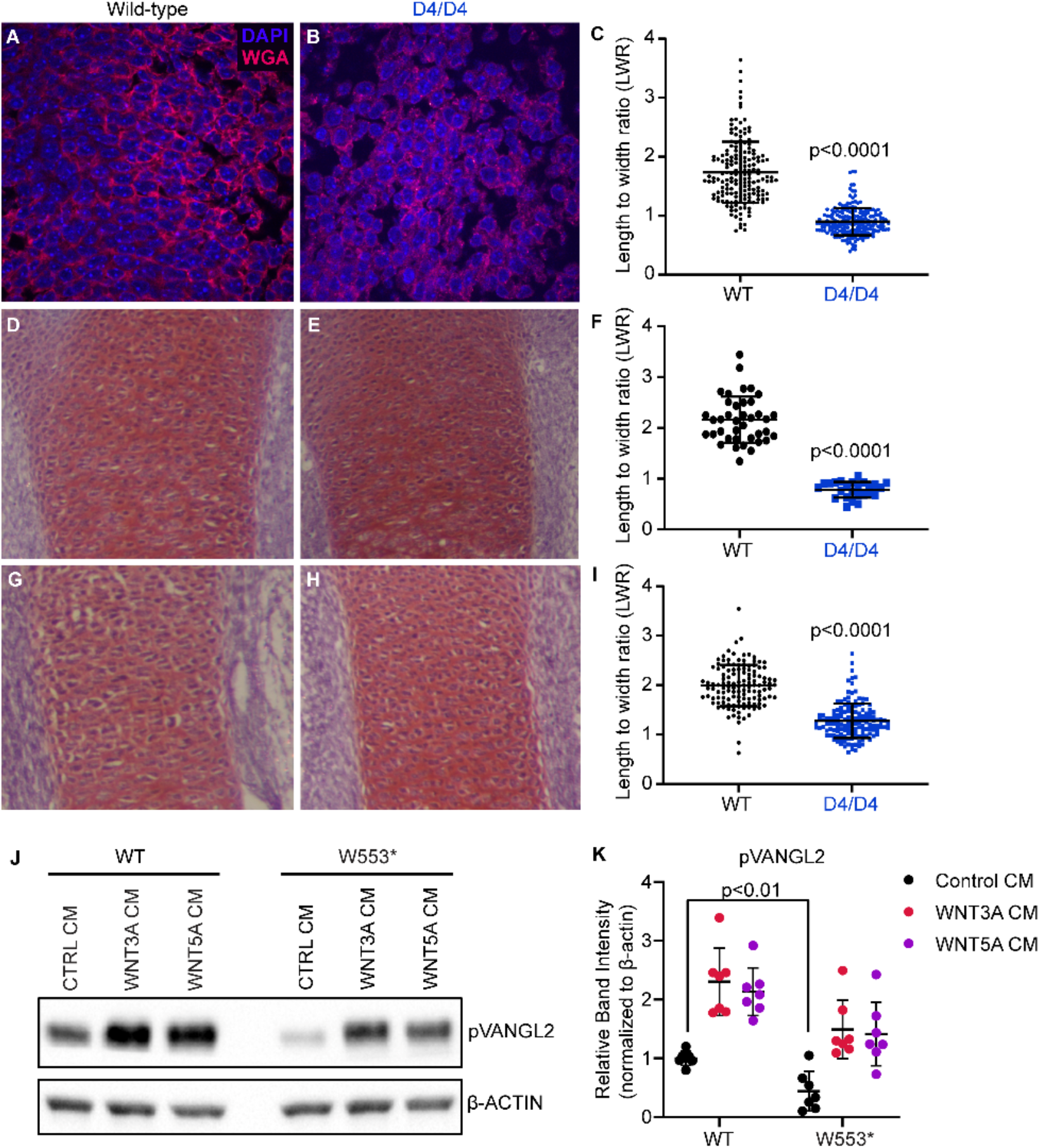
Altered morphology of *Fzd2*^*em1Rstot(D4)*^ limb chondrocytes. (**A** and **B**) Representative images of sagittal sections of E17.5 wild-type and *Fzd2*^*em1Rstot(D4)*^ homozygote femur stained for wheat germ agglutinin (WGA, red) and DAPI (blue). (**C**) Chondrocyte width and length were measured in the midshaft of the femur. Each dot represents one chondrocyte measured from 2 WT and 2 D4/D4 embryos for a total of n=277 WT chondrocytes and n=153 D4/D4 chondrocytes. (**D** and **E**) Representative images of sagittal sections of E17.5 wild-type and *Fzd2*^*em1Rstot(D4)*^ homozygote femur stained for safranin-O. (**F**) Chondrocyte width and length were measured in the midshaft of the femur. Each dot represents one chondrocyte measured from 3 WT and 3 D4/D4 embryos for a total of n=39 WT chondrocytes and n=25 D4/D4 chondrocytes. (**G** and **H**) Representative images of sagittal sections of E17.5 wild-type and *Fzd2*^*em1Rstot(D4)*^ homozygote forelimb stained for safranin-O. (**I**) Chondrocyte width and length were measured in the midshaft of the forelimb. Each dot represents one chondrocyte measured from 3 WT and 3 D4/D4 embryos for a total of n=128 WT chondrocytes and n=141 D4/D4 chondrocytes. (**J-K**) Mouse embryonic fibroblasts (MEFs) were generated from WT or W553* embryos and cultured with LGK-974 (1 µM) for 48 hours followed by conditioned media treatment (1/5 vol:vol) for 24 hours. Immunoblot analysis for pVANGL2 was performed and normalized to β-ACTIN. n=7/group. All p values are indicated on graphs.

### Limb phenotypes were rescued in *Fzd2*^*D4/D4*^ embryos by stimulating canonical signaling

A previous study had demonstrated the power of modulating the Wnt pathway in craniofacial development to rescue the cleft palate in *Pax9* mutant mice (29). We reasoned that the action of the IIIC3a molecule as an antagonist of the Wnt antagonist DKK may act to augment the pathway and partially rescue the genetic lesions in *Fzd2*. We tested the activity of IIIC3a *in vitro* and found that IIIC3a increased the canonical Wnt signaling reporter TOPFlash in a dose-dependent manner in the presence of WNT3A conditioned media (**Supplemental Figure 3A-B**). We tested this hypothesis with the *Fzd2*^*D4*^ allele and treated pregnant dams with daily 25 mg/kg intraperitoneal injections of IIIC3a from E10.5 to E14.5 and examined embryos at E17.5 (**Figure 9A**).

**Figure 9.**
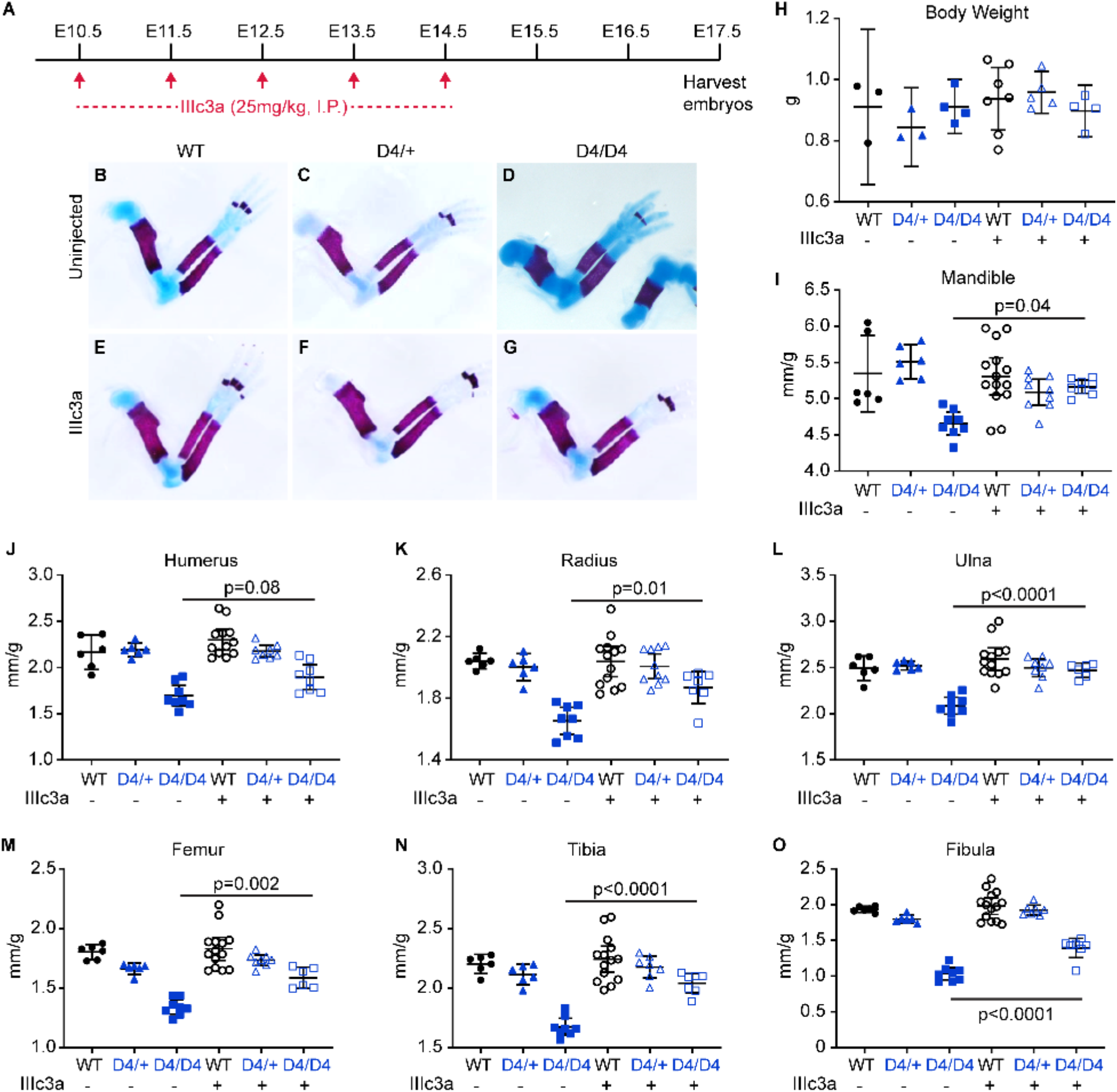
Inhibition of Dkk1 *in utero* normalizes limb lengths in *Fzd2*^*em1Rstot(D4)*^ homozygotes. (**A**) IIIC3a was administered daily via intraperitoneal injection to pregnant dams from E10.5-E14.5 and embryos were harvest at E17.5. (**B-G**) Whole mount images of skeletal preps of forelimbs from E17.5 wild-type, heterozygous and homozygous D4 animals treated with or without IIIC3a (25 mg/kg). (**H**) Body weight measurements. n=3-5/group. (**I**) Mandible length (n=2 bones per animal) per body weight. (**J-O**) Length measurements of humeri, radii, ulnae, femora, tibiae, and fibulae (n=2 per animal) per embryo body weight. All p values are indicated on graphs

The body weight of the embryos did not differ significantly (**Figure 9H**) and all *Fzd2*^*D4/D4*^ homozygotes (n=4) still had cleft palate. We measured the long bones and mandible and normalized to body weight (**Figure 9I-O**). We found that all bones measured in IIIC3a-treated *Fzd2*^*D4/D4*^ homozygotes were significantly longer than untreated *Fzd2*^*D4/D4*^ homozygotes and very similar to all control genotypes. Thus, we conclude that although both canonical and non-canonical Wnt signaling are affected, modulating the canonical Wnt pathway was able to significantly rescue long bone length.

## DISCUSSION

In this study, we found that disruptions of the C-terminal region of FZD2 in mice leads to highly penetrant and severe craniofacial abnormalities resulting in pre-weaning lethality. Disruption of the DVL-binding domain (22, 30, 31), even a one amino-acid deletion (*Fzd2*^*D3*^), was sufficient to lead to autosomal recessive cleft palate. More substantial disruption of the DVL-binding domain via a frameshift in this region (*Fzd2*^*D4*^) leads to a similar phenotype with higher penetrance of cleft palate in *Fzd2*^*D4/D4*^ mice, as well as an incompletely penetrant dominant failure to thrive in *Fzd2*^*D4/+*^ animals. Intriguingly, the most severe phenotypes were associated with the ortholog of the variant identified in the human genetics studies, *Fzd2*^*W553\**^. For this allele we were unable to generate live heterozygous or homozygous mice at P14; however, via a series of independently generated “F0” CRISPR-modified mice we were able to identify embryos with mosaic *Fzd2*^*W553\**^ knock-in. Such *Fzd2*^*W553\**^-edited embryos all exhibited cleft palate, decreased weights, and significantly shorter long-bone lengths as compared to unedited littermates. This clearly demonstrates a dominant inheritance mechanism for *Fzd2*^*W553\**^ and recapitulates virtually all the phenotypes observed in the human FZD2-associated AD-RS we previously attributed to this variant.

The mechanism of the incompletely penetrant failure to thrive and ultimate cause of death in non-cleft *Fzd2*^*D4/+*^and *Fzd2*^*D3/D3*^ animals remains unclear. *Fzd2*^*D4/+*^ pups are born in normal Mendelian ratios, although it is clear by 1-2 weeks of age that there is a subset of mice that are substantially smaller than their littermates. This failure-to-thrive was also seen previously in non-cleft *Fzd2*^*null*^ mice where a thorough investigation again failed to yield a cause of death (19).

The mechanism of dominance resulting from C-terminal disruption of FZD2 also remains unclear but may be due to the fact that the C-terminal intracellular domain of FZD2 is disrupted while the sequence encoding the Wnt-binding extracellular domains of FZD2 remains intact. We hypothesize that if Wnt ligands can bind the C-terminal FZD2 mutants but are unable to mediate a downstream signaling cascade, FZD2 may act as a “sink” for Wnt signaling components and disrupt signaling of other *Fzd* family members through their overlapping expression patterns and interactions with *Wnt* ligands that bind multiple frizzled proteins. There is also evidence that Fzd receptors form multimers in the membrane (32, 33). In this way, one FZD2 protein with a C-terminal modification may ultimately affect the signaling capacity of many more proteins and produce a “dominant-negative” effect. Additional studies are necessary to identify biochemical changes in FZD2-DVL binding interactions as well as the effects of FZD2-signaling disruption on other members of the WNT-FZD signaling pathway.

Our findings that both canonical and non-canonical Wnt signaling are affected by these *Fzd2* variants suggested that pharmacological intervention targeted at the canonical arm of the pathway might ameliorate some of the phenotypes. Indeed, we saw that IIIC3a addition was able to restore more normal length to the long bones, but the cleft palate incidence was refractory to this intervention. This suggests that non-canonical effects on convergent extension during craniofacial development may be too profound for this treatment or that alternate doses should be attempted. Regardless of that potential result, the long bone deficits are much more difficult to treat in these patients than surgical repair of the cleft palate. A medical intervention would appear to be a very attractive option for therapeutic treatment, especially in a case where the genetic diagnosis could potentially be made years before long-bone growth is concluded in human development.

Overall, this study demonstrates an essential role for the signaling functions of the C-terminal region of FZD2 and validates the findings of FZD2-associated AD-RS we previously observed in human patients. This work also validates the approach of CRISPR/Cas9-mediated knock-in of human variants as an important tool for investigating presumably causative sequence variants. Importantly, the use of the *i-*GONAD technique allows for the rapid production of these alleles and decreases the number of animals needed to generate embryos harboring variants of interest. As the availability of human patient exome/genome sequencing becomes higher due to ever-decreasing costs, many previously unreported rare variants will be identified. While large-scale mouse knockout projects provide valuable information as to the function of understudied genes, most human sequence variation is not comprised of highly-orchestrated deletions of entire coding sequences leading to fully null alleles, but rather of small single nucleotide variants (34-36). As such, there is tremendous value in being able to study specific patient variants encoding either hypomorphic alleles or gain-of-function alleles that would not be adequately modeled by a complete loss of function. For some genes in which loss-of-function is well tolerated, studying such gain-of-function alleles may, in fact, be the most efficient way to increase our understanding of the normal function of a particular gene.

## METHODS

### Animal housing

All experiments utilizing mice in this study were performed utilizing ethically acceptable procedures as approved by the Institutional Animal Care and Use Committee at Cincinnati Children’s Hospital Medical Center and Van Andel Institute.

Mice were fed LabDiet 5021 mouse breeder diet and housed in ventilated cages with a 12-hour light/12-hour dark cycle.

### Identifying sgRNA sequences

Identification of optimal sgRNA targets was performed using the MIT genome engineering tool (http://crispr.mit.edu). The top three guide sequences were cloned into PX458M; MK4 cells cultured under standard conditions (DMEM + 10% FBS + 1% P/S) were then transfected with each plasmid using Lipofectamine 3000 per the manufacturer’s protocol (Invitrogen). After 24h, genomic DNA was isolated using NaOH lysis of cells. A region flanking the target region was PCR-amplified, and the Surveyor mutation detection kit (IDT) was used per manufacturer’s instructions to digest the reannealed PCR product. The cleaved products were separated on an agarose gel which was quantified via Gel-Imager for PC. The sgRNA showing highest levels of editing was utilized for all subsequent experiments.

### Zygote microinjection/generation of edited embryos

Mouse zygotes (C57BL6/N strain) were injected with 200 ng/µL Cas9 protein (IDT and ThermoFisher), 100 ng/µL Fzd2-specific sgRNA (GCAAGACACTGCACTCGTGG), and 75 ng/µL single-stranded donor oligonucleotide (GTGGGCATCACGTCGGGCTTCTGGATCTGGTCCGGAAAAACTCTTCATTCTTGATGATAGTTCTACACTCGTCTCACC AACAGCCGGCATGGCGAGACCACTGTGTGAAGC; IDT, Iowa) followed by surgical implantation into pseudo-pregnant female (CD-1 strain) mice. Two rounds of microinjection were performed. The resulting live born pups from the first round were weaned and used for additional screening and mating. Pregnant females from the second round were euthanized at E17.5 and embryos were screened for phenotypic changes and genetic modifications.

### *i*mproved Genome Editing of Oviductal Nucleic Acid Delivery (*i-*GONAD)

To rapidly generate additional embryos with the *Fzd2*^*W553\**^ variant, the *i-*GONAD technique was used (23, 24). The night before surgery, B6C3F1/J (JAX stock #100010) males (8-16 week old) and females (7-12 week old) were bred. Vaginal plugs were detected the following morning at 9AM. Complete sgRNA’s from IDT (CCGGCAAGACACTGCACTCG; crRNA and tracrRNA) were incubated with Cas9-mSA (37) protein to generate ribonucleoprotein (RNP) complexes for 10 minutes at room temperature before the biotinylated template (ACATGATCAAATACCTCATGACGCTCATCGTGGGCATCACGTCGGGCTTCTGGATCTGGTCCGGCAAGACACTGCATTCATGAAGGAAGTTCTACACTCGTCTCACCAACAGCCGGCATGGCGAGACCACTGTGTGAAGCGGTCTCGCCTGCCTGCCGGGCTT; IDT) was added (38). The final *i-*GONAD mix contained 30 µM sgRNA: 0.6 µg/µL Cas9-mSA protein, 2 µg/µl BIO-template (in filtered 1 mM Tris-HCl, pH 7.5; 0.1 mM EDTA). Females which were identified with vaginal plugs underwent the *i-*GONAD technique at approximately E0.7 (4:00PM). Anesthesia (1X Avertin, intraperitoneal, ∼45 units per 20-gram mouse) and analgesia (5 mg/kg carprofen, subcutaneous) were administered, and deep anesthesia tested via toe pinch reflex. Under a surgical microscope, the mouse was placed dorsal side up, sprayed with 70% ethanol and wiped with a Kimwipe. A 10 mm incision was made just inferior to the ribs, through the skin centrally, and sprayed with 70% ethanol and wiped with a Kimwipe. The incision was adjusted to expose the right or left flank and a 5-mm incision was made through the muscle layer. A sterile surgical drape was placed over the incision and the ovary was pulled out of the body cavity by grasping the fat pad. The fat pad was then clamped with an Aorta-Klemme to stabilize the tissue.

Approximately 1.5 µL of appropriate CRISPR reagents were administered via mouth pipette to the oviduct just upstream of the ampulla containing the fertilized eggs. A 5 mm x 5 mm Kimwipe strip soaked in 1X PBS was wrapped around the injected oviduct and the oviduct was placed firmly within the probes of the electrokinetic tweezers (NepaGene, CUY652P2.5 x4) and electroporated with 8 pulses at 50V, 5 msec per pulse and 1 second interval in between pulses (BTX T820 square wave electroporator). Following electroporation, the Kimwipe was removed, and the ovary placed back into the body cavity. The procedure was repeated on the other ovary/oviduct and the skin incision closed with two surgical staples. Animals were placed back in their cage on a warmer and monitored until completely recovered.

### Genotyping by PCR and indel sequencing

The resulting embryos or live born pups were screened for evidence of editing via PCR/Sanger sequencing of embryonic yolk sacs or tail clips. Genomic DNA was amplified using forward GTAAGCCAGCACTGCAAGAG and reverse GTGAAGGAGGGCACGGTG primers. Pups exhibiting editing of interest were then crossed to WT (C57BL6/J) mice and the resulting progeny were Sanger sequenced (CCHMC DNA Sequencing and Genotyping Core) to confirm the alleles generated. Sanger sequences were analyzed via “4-Peaks” software (Mac). In-silico analysis (Benchling.com) was utilized to analyze the consequences of each allele on the predicted putative proteins.

### Genotyping by Next Generation Sequencing

*Fzd2*^*W553*^*\** animals generated via *i-*GONAD were genotyped via Next Generation Sequencing. Briefly, the targeted region (including ∼250bp regions at both ends) was PCR amplified with primers including unique 8 bp barcodes per sample (**Supplemental Table 2**). The PCR products were run on gel to confirm the purity and pooled before being sent for Amplicon EZ Sequencing (Genewiz/Azenta, please see the vendor’s website for more technical details). The reads were then aligned to the wildtype Fzd2 DNA sequence with the aligner BWA-MEM. After demultiplexing the reads based on the barcodes, reads were visualized with the software IGV (BROAD Institute) to identify the modifications at the target region.

### “F0” embryonic analyses

Embryos generated via the second round of microinjection or *i-*GONAD were collected at E16.5-E18.5 and analyzed for cleft palate. Embryos were imaged in PBS prior to preparing tissue for downstream histological assays, skeletal preparations, or micro-CT analyses.

### Phenotypic characterization of Fzd2 D3/D4 alleles

To identify potential survival defects in CRISPR-generated alleles, a het x het breeding scheme was utilized. Resulting pups were genotyped via PCR followed by either restriction enzyme digest/agarose gel electrophoresis or Sanger sequencing.

Survival vs. expected Mendelian ratios were scored at late embryonic stages (E15.5-E18.5) weaning age (P18-P28) and analyzed for significance via Student’s t-test (GraphPad Prism) with p<0.05 as a threshold for significance. Examination and imaging of embryonic phenotypes was performed via whole mount imaging in PBS, histological analyses and skeletal preparations.

### Histology

H&E fixation in Bouin’s solution followed by washes in 70% ethanol, and H&E staining. For H&E staining, embryos were embedded in paraffin and cut to 10 µm sections before staining using standard techniques (39). All images were taken via Zeiss Discovery.V8 Stereoscope. Paired images are shown at the same magnification.

### Palate proliferation & apoptosis

Embryos were collected at E13.5, fixed in 4% PFA for 2 h followed by cryoprotection in 30% sucrose and embedding in OCT compound. 10 µm frozen sections were cut and transferred to glass slides. Antibody staining for phospho-histone H3 and cleaved-caspase 3 was performed using standard immunohistochemistry techniques. Briefly, sections were blocked in 4% Normal Goat Serum (NGS) for 1 h, incubated overnight with Cleaved Caspase-3 (CC-3; Cell Signaling 9661S; 1:300) or phospho-Histone H3 (pHH3; Sigma H0412; 1:500) diluted in blocking buffer. Following washing with PBS, slides were incubated for 1 h with Alexa Fluor® 488 Goat Anti-Rabbit IgG (Invitrogen A11008: 1:500) diluted in blocking buffer. Sections were briefly stained with DAPI and coverslipped. Images were taken on a Nikon C2 confocal microscope. Paired images are shown at the same magnification.

### Wheat germ agglutinin (WGA) staining

Embryos were collected at E12.5, fixed in 4% PFA overnight, and forelimbs embedded in OCT (Sakura) prior to cryosectioning. Samples were sagittal sectioned (10 µm) and stained for WGA (5 µg/mL in HBSS) for 3 min at room temperature. Slides were washed in PBS followed by staining with DAPI (Thermofisher 1 ug/ml) and mounting with ProLong Gold (Invitrogen). Paired wild-type and *Fzd2*^*em1Rstot(D4)*^ homozygote images were taken at the same magnification using a Nikon C2 confocal microscope. Length to width ratios were calculated for chondrocytes from the midshaft of the forelimb from 3 *Fzd2*^*+*^ and 3 *Fzd2*^*em1Rstot(D4)*^ homozygote embryos.

### Safranin O staining

Femur and ulna from E12.5 were fixed in 4% PFA and processed using standard histology techniques. Sections (10 µm) were stained for safranin O following a previously published protocol with modifications (40). Briefly, slides were deparaffinized, washed in dH2O, and stained with hematoxylin (Hematoxylin Solution, Harris Modified, Millipore Sigma) for 5 min. Slides were washed with tap water followed by 5 dips in acid ethanol (1:400 HCl in 70% ethanol) and washed again with tap water. Slides were then stained with 0.001% Fast Green FCF (C.I. 42053) for 5 min followed by 1% acetic acid for 10-15 s and 0.1% Safranin O for 5 min. Slides were then dehydrated and paired wild-type and *Fzd2*^*em1Rstot(D4)*^ homozygote images were taken at the same magnification using a Zeiss Discovery.V8 Stereoscope. Length to width ratios were calculated for chondrocytes from the midshaft of the forelimb or femur from 3 WT and 3 *Fzd2*^*em1Rstot(D4)*^ homozygote embryos.

### Genetic background effects

*Fzd2* D4/+ (C57BL/6J; Jackson Lab, strain #000664) background animals were crossed with WT CD-1 mice (Charles River, strain #022). Heterozygous D4/+ males and females from the F1 generation (50% B6, 50% CD-1) were crossed to produce an F2 generation that was phenotypically evaluated and genotyped to identify survival defects at both late embryonic time points (E15.5-E18.5) and at weaning (P18-P28).

### Skeletal preparations

Skeletons from E17.5 animals were stained for alcian blue and alizarin red to visualize cartilage and bone, respectively. Briefly, embryos were eviscerated and fixed for 2 days in 95% ethanol. They were stained overnight at room temperature in 0.03% (w/v) Alcian Blue solution (Sigma-Aldrich, A3157) containing 80% ethanol and 20% glacial acetic acid. Samples were destained in 95% ethanol for 24 h followed by pre-clearing in 1% KOH overnight at room temperature.

Skeletons were then stained overnight in 0.005% Alizarin Red solution (Sigma-Aldrich, cat#A5533) containing 1% KOH. A second round of clearing was performed by incubating tissues in 20% glycerol/1% KOH solution for 24 h. Finally, they were transferred to 50% glycerol/50% ethanol for photography. Skeletal preparations were imaged using a Zeiss Discovery.V8 Stereoscope and length measurements were recorded for mandibular bones, humeri, ulnae, radii, femora, tibiae, and fibulae.

### Micro Computed Tomography

Skulls and limbs from E18.5 *Fzd2*^*W553\**^ animals were examined using the SkyScan 1172 micro-computed tomography (µCT) system (Bruker MicroCT: Kontich, Belgium). Tissues were fixed in 10% neutral buffered formalin (NBF) at room temperature for 48 h, stored in 70% ethanol, and kept in the 70% ethanol solution for scanning.

Specimens were scanned using an X-ray voltage of 50 kV, current of 201 µA, and 0.5 mm aluminum filter. The samples were imaged using 2000 × 1200 pixel resolution and 8 µm image pixel size and reconstructed using NRecon 1.7.4.6 (Bruker MicroCT).

For each sample, the volume of interest (VOI) was defined using DataViewer 1.5.6.3 (Bruker MicroCT) and and a region of interest (ROI) was defined using CTAn 1.18.8.0 (Bruker MicroCT). The defined ROI was used to generate a 3d surface rendered model in CTAn which was then visualized and manipulated in CTVol 2.2.1.0 (Bruker MicroCT).

### Quantitative real time PCR (qRT-PCR)

qRT-PCR was performed using standard techniques. RNA was extracted from craniofacial tissues of E13.5 *Fzd2*^*+*^ and *Fzd2*^*D4/D4*^ littermate embryos using Qiagen RNA extraction kit. RNA was reverse transcribed using Superscript III First-Strand Synthesis System for RT-PCR (Invitrogen), and cDNA was amplified and detected using TaqMan Universal PCR master mix (Applied Biosystems) and TaqMan probes, including mouse Axin2 (Mm00443610_m1), mouse Dkk1 (Mm00438422_m1) and mouse Gapdh (Mm99999915_g1). Realtime PCR was analyzed on Applied Biosystems QuantStudio 6 (ThermoFisher).

### IIIC3a in vitro activity

HEK293-Super TOPFlash (STF) (gift from Jeremy Nathans (41)) or C3H10T1/2-STF (ATCC CCL-226) cells were treated with 1 µM LGK-974 (Cayman Chemical 14072) for 48 h to inhibit endogenous WNT secretion. Dkk Inhibitor II (IIIC3a; Calbiochem; EMD_BIO-317701) was added at varying concentrations (0-250 µM) with CTRL CM (1/10 dilution) or WNT3A CM (1/10 dilution) made from L cells (ATCC L Cells CRL-2648, ATCC L-Wnt-3A CRL-2647, ATCC L Wnt-5A CRL-2814).

Transactivation of a β-catenin/TCF-responsive reporter construct was measured using the Promega Dual-Luciferase™ Reporter (DLR™) Assay to identify activated canonical WNT signaling. Each treatment had 4 technical replicates and the experiments was performed twice with consistent results.

### IIIC3a treatment

Dkk Inhibitor II (IIIC3a; Calbiochem; EMD_BIO-317701) was administered (25 mg/kg; intraperitoneal) to pregnant females on E10.5, E11.5, E12.5, E13.5 and E14.5. A stock solution of IIIC3a was made at 25 mg/mL in DMSO and diluted 1:10 in 1X dPBS (2.5 mg/mL) for working solution on the day of injections.

### Mouse embryonic fibroblasts (MEFs)

Following *i*-GONAD surgery to induce the W553* variant in mice, E12.5 embryos were harvested to generate MEFs according to standard protocols (42), and yolk sacs were collected for Sanger sequencing and next generation sequencing. Briefly, E12.5 embryos were minced in 0.05% Tryspin-EDTA (Gibco) and incubated for 30 m at 37 °C. Cells were then neutralized with DMEM with 10% fetal bovine serum (FBS) and 1% penicillin/streptomycin and cultured for 48 h in T75 flasks. Cells used for experiments were passage 3 or fewer. To test noncanonical signaling changes in MEFs, cells were first treated with LGK-974 (1 µM) for 48 hours to inhibit endogenous WNT ligand secretion. MEFs were then serum starved for 24 hours (with LGK-974) followed by a 24 hour treatment with CM made from L cells. CM was diluted 1/5 in DMEM with 10% FBS. Cells were rinsed once with cold 1X PBS and lysed with lysis buffer (50mM Na_2_HPO_4_, 1mM sodium pyrophosphate, 20mM NaF, 2mM EDTA, 2mM EGTA, 10mM NaCl, 1% TritonX-100, 1mM DTT, and cOmplete™ Mini EDTA-free Protease Inhibitor Cocktail (Roche)).

Cells were centrifuged at 21000 x g for 20 minutes at 4 °C and the supernatant collected. Samples were resolved on Mini-PROTEAN TGX Stain-Free Gels (BIO-RAD, 456026) and transferred using the Trans-Blot Turbo transfer system (BIO-RAD). The membranes were blocked with EveryBlot Blocking Buffer (BIO-RAD, 12010020) and probed with pVANGL2 (Invitrogen, MA5-38242, 1:1000) antibody followed by Anti-Mouse IgG HRP-linked antibody (Cell Signaling Technologies, 7074, 1:1000) and chemiluminescent visualization on a BIO-RAD ChemiDoc.

### Statistics

All statistical analyses were performed using GraphPad Prism 9.3.1. Ordinary one-way ANOVA with Dunnett’s post hoc multiple comparisons test was performed for comparison of D3/D4 heterozygotes and homozygotes to wild-type samples. Ordinary one-way ANOVA with Tukey’s post hoc multiple comparisons test was performed for IIIC3a treatment studies. Unpaired t test was performed for comparison of W5553* heterozygotes or D4 homozygotes to wild-type samples. P values are indicated on all graphs. The data presented are mean +/- 95% confidence interval.

### Study approval

All experiments utilizing mice in this study were performed utilizing ethically acceptable procedures as approved by the Institutional Animal Care and Use Committees at Cincinnati Children’s Hospital Medical Center and Van Andel Institute.

## Author contributions

RWS, RPL, MNM, BOW designed research studies; RPL, MNM, SV, EB, EF, CAM, conducted experiments; RPL, MNM, SV, CRD, ZZ, BOW, RWS analyzed data; RPL, MNM, RWS wrote the original draft of the manuscript. All authors reviewed and edited the manuscript. The order of the co–first authors was determined by a thorough discussion of the relative contributions to experimental design, execution, and manuscript writing. The authors fully endorse the right of the co-first authors to rearrange the first three authors to suit their needs.

## ACKNOWLEDGEMENTS

We thank the Cincinnati Children’s Hospital (CCHMC) Transgenic Core for assistance with zygote injections and the VAI Vivarium and Transgenic Core staff for assistance with the *i-*GONAD injections. This work is supported by the NIH grants DE027091 to RWS and DE031039 to MNM. Additional funds for this work were provided by the CCHMC Center for Pediatric Genomics, the Cincinnati Children’s Research foundation, and the Van Andel Institute.

