## Supplemental Material for "Dkk1 inhibition normalizes limb phenotypes in a mouse model of *Fzd2* associated omodysplasia Robinow syndromes"

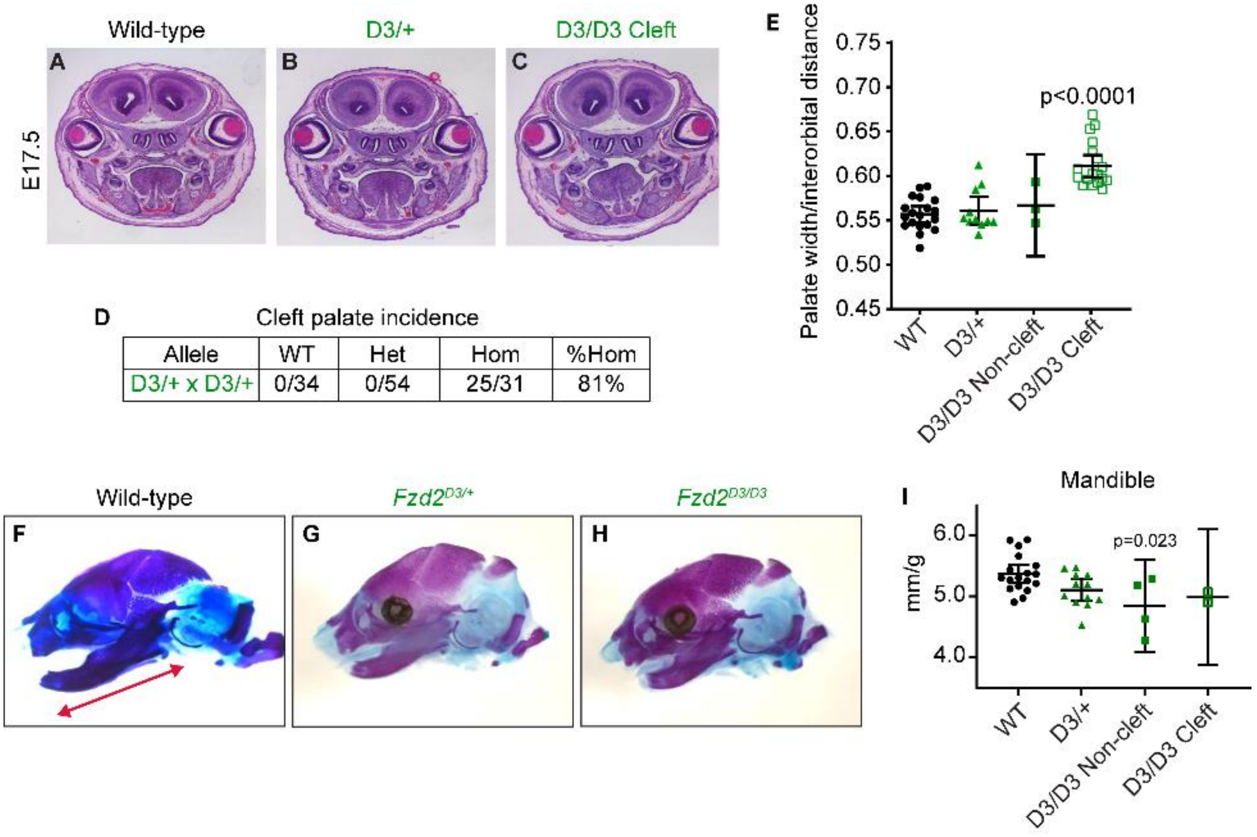


**Supplemental Figure 1.** ***$Fzd2^{em1Rstot(D3)}$*** **homozygotes have partially penetrant cleft palate.** (**A-C**) Coronal sections through the mid-palate of E17.5 wild-type, heterozygous and homozygous D3 animals. (**D**) Incidence of cleft palate in E17.5 embryos recovered from D3/+ x D3/+ matings. (**E**) Palatal width as a percentage of interorbital width. n=9-21/group. (**F-H**) Left side view images of skeletal preps of E17.5 wild-type, heterozygous and homozygous D4 skulls. Red double-ended arrow in panel **F** indicates the plane in which the mandible was measured. (**I**) Mandible length (n=2 bones per animal) per body weight.

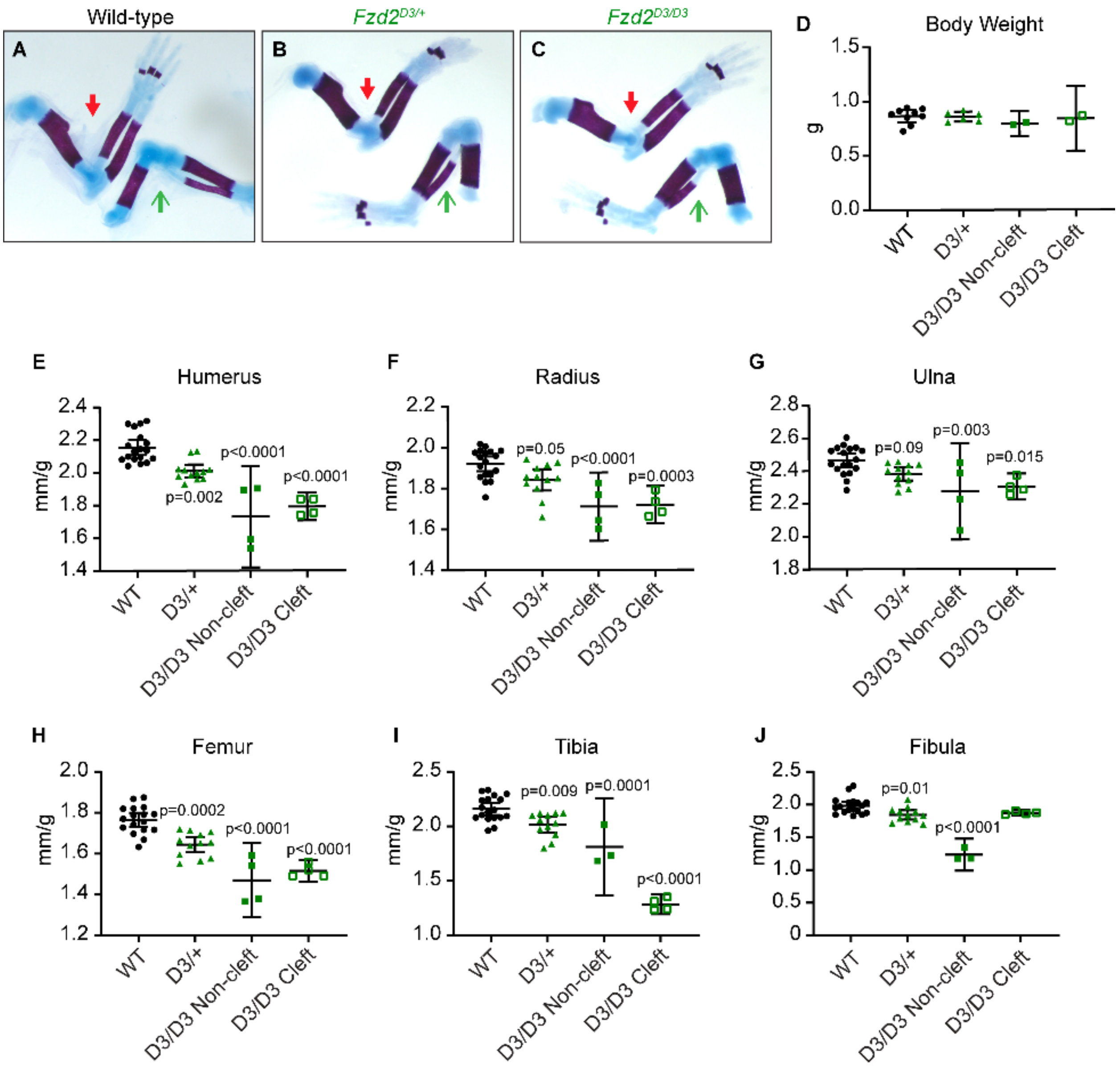


**Supplemental Figure 2. Craniofacial and limb elements are shorter in E17.5 *Fzd2*$^{em1Rstot(D3)}$ homozygotes.** (**A-C**) Whole mount images of skeletal preps of forelimbs (red closed arrows) and hindlimbs (green open arrows) from E17.5 wild-type, heterozygous and homozygous D3 animals. (**D**) Body weight measurements. n=3-4/group. (**E-J**) Length measurements of humeri, radii, ulnae, femora, tibiae, and fibulae (n=2 per animal) per embryo body weight. All p values are indicated on graphs.

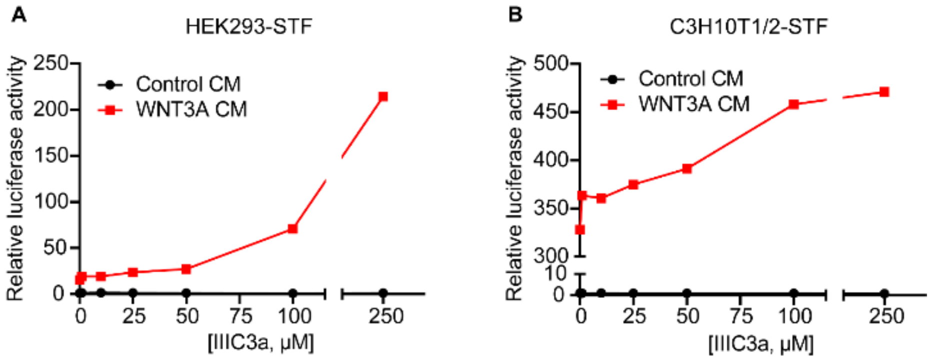


**Supplemental Figure 3. IIIC3a activity *in vitro*.** (**A**) 293-STF cells were treated with varying concentrations of DKK inhibitor, IIIC3a in conjunction with CTRL CM (1/10 dilution) or WNT3A CM (1/10 dilution). STF luciferase activity was measured and normalized to control conditioned media treated cells. n=4 technical replicates/treatment. (**B**) C3H10T1/2-STF were treated with varying concentrations of Dkk inhibitor, IIIC3a in conjunction with CTRL CM (1/10 dilution) or WNT3A CM (1/10 dilution). Luciferase activity was measured. n=4 technical replicates/treatment.

**Supplemental Table 1. Mouse FZD2 C-terminal modifications resulting in cleft palate.**

| **Gene** | **Variant type** | **Zygosity** | **CDS variant** | **Amino Acid variant** | **Cleft palate** |
|---|---|---|---|---|---|
| *Fzd2* | nonframeshift insertion | mosaic | c.1653_1654insTTT | p.His551_Ser552insPhe | No |
| *Fzd2* | -1 frameshift | mosaic | c.1653del | p.Ser552Argfs*62 | Yes |
| *Fzd2* | nonframeshift deletion | compound het | c.1650_1655del<br>c.1654_1656del | p.His551_Ser552del<br>p.Ser552del | Yes |
| *Fzd2* | -1 frameshift | mosaic | c.1645_1669del | p.Ser552Thrfs*57 | Yes |
| *Fzd2* | +1 frameshift | mosaic | c.1653delCinsAA | p.His551Glnfs*59 | Yes |
| *Fzd2* | -1 frameshift, nonframeshift deletion | compound het | c.1648_1654del<br>c.1652_1654del | p.Leu550Argfs*62<br>p.His551Pro | Yes |
| *Fzd2* | -1 frameshift | mosaic | c.1653delCinsGGAAGT | p.His551Glnfs*65 | Yes |

**Supplemental Table 2. Barcoded primers for next generation sequencing.**

| *Barcode Sequence only* | *Forward Primer Sequence only* | *Forward Primer Sequence* | *Reverse Primer Sequence only* | *Reverse Primer Sequence* |
|---|---|---|---|---|
| cggttcaa | TAAGCCAGCACTGCAAGAG | cggttcaaTAAGCCAGCACTGCAAGAG | GCCCTGGTGTCTTCGATTT | cggttcaaGCCCTGGTGTCTTCGATTT |
| gctggatt | TAAGCCAGCACTGCAAGAG | gctggattTAAGCCAGCACTGCAAGAG | GCCCTGGTGTCTTCGATTT | gctggattGCCCTGGTGTCTTCGATTT |
| taactcgg | TAAGCCAGCACTGCAAGAG | taactcggTAAGCCAGCACTGCAAGAG | GCCCTGGTGTCTTCGATTT | taactcggGCCCTGGTGTCTTCGATTT |
| taacagtt | TAAGCCAGCACTGCAAGAG | taacagttTAAGCCAGCACTGCAAGAG | GCCCTGGTGTCTTCGATTT | taacagttGCCCTGGTGTCTTCGATTT |
| atactcaa | TAAGCCAGCACTGCAAGAG | atactcaaTAAGCCAGCACTGCAAGAG | GCCCTGGTGTCTTCGATTT | atactcaaGCCCTGGTGTCTTCGATTT |
| gctgagaa | TAAGCCAGCACTGCAAGAG | gctgagaaTAAGCCAGCACTGCAAGAG | GCCCTGGTGTCTTCGATTT | gctgagaaGCCCTGGTGTCTTCGATTT |
| attggagg | TAAGCCAGCACTGCAAGAG | attggaggTAAGCCAGCACTGCAAGAG | GCCCTGGTGTCTTCGATTT | attggaggGCCCTGGTGTCTTCGATTT |
| tagtctaa | TAAGCCAGCACTGCAAGAG | tagtctaaTAAGCCAGCACTGCAAGAG | GCCCTGGTGTCTTCGATTT | tagtctaaGCCCTGGTGTCTTCGATTT |
| cggtgacc | TAAGCCAGCACTGCAAGAG | cggtgaccTAAGCCAGCACTGCAAGAG | GCCCTGGTGTCTTCGATTT | cggtgaccGCCCTGGTGTCTTCGATTT |
| tacagagg | TAAGCCAGCACTGCAAGAG | tacagaggTAAGCCAGCACTGCAAGAG | GCCCTGGTGTCTTCGATTT | tacagaggGCCCTGGTGTCTTCGATTT |
| attgtcaa | TAAGCCAGCACTGCAAGAG | attgtcaaTAAGCCAGCACTGCAAGAG | GCCCTGGTGTCTTCGATTT | attgtcaaGCCCTGGTGTCTTCGATTT |
| tatgtctt | TAAGCCAGCACTGCAAGAG | tatgtcttTAAGCCAGCACTGCAAGAG | GCCCTGGTGTCTTCGATTT | tatgtcttGCCCTGGTGTCTTCGATTT |
| attggatt | TAAGCCAGCACTGCAAGAG | attggattTAAGCCAGCACTGCAAGAG | GCCCTGGTGTCTTCGATTT | attggattGCCCTGGTGTCTTCGATTT |
| atactcgg | TAAGCCAGCACTGCAAGAG | atactcggTAAGCCAGCACTGCAAGAG | GCCCTGGTGTCTTCGATTT | atactcggGCCCTGGTGTCTTCGATTT |
| tatgagaa | TAAGCCAGCACTGCAAGAG | tatgagaaTAAGCCAGCACTGCAAGAG | GCCCTGGTGTCTTCGATTT | tatgagaaGCCCTGGTGTCTTCGATTT |
| gcacagtt | TAAGCCAGCACTGCAAGAG | gcacagttTAAGCCAGCACTGCAAGAG | GCCCTGGTGTCTTCGATTT | gcacagttGCCCTGGTGTCTTCGATTT |
| cgtggatt | TAAGCCAGCACTGCAAGAG | cgtggattTAAGCCAGCACTGCAAGAG | GCCCTGGTGTCTTCGATTT | cgtggattGCCCTGGTGTCTTCGATTT |
| tagtagaa | TAAGCCAGCACTGCAAGAG | tagtagaaTAAGCCAGCACTGCAAGAG | GCCCTGGTGTCTTCGATTT | tagtagaaGCCCTGGTGTCTTCGATTT |
| gcacgatt | TAAGCCAGCACTGCAAGAG | gcacgattTAAGCCAGCACTGCAAGAG | GCCCTGGTGTCTTCGATTT | gcacgattGCCCTGGTGTCTTCGATTT |

| | | | | |
|---|---|---|---|---|
| cggtagcc | TAAGCCAGCACTGCAAGAG | cggtagccTAAGCCAGCACTGCAAGAG | GCCCTGGTGTCTTCGATTT | cggtagccGCCCTGGTGTCTTCGATTT |
| tagttctt | TAAGCCAGCACTGCAAGAG | tagttcttTAAGCCAGCACTGCAAGAG | GCCCTGGTGTCTTCGATTT | tagttcttGCCCTGGTGTCTTCGATTT |
| tacaagtt | TAAGCCAGCACTGCAAGAG | tacaagttTAAGCCAGCACTGCAAGAG | GCCCTGGTGTCTTCGATTT | tacaagttGCCCTGGTGTCTTCGATTT |
| atcactgg | TAAGCCAGCACTGCAAGAG | atcactggTAAGCCAGCACTGCAAGAG | GCCCTGGTGTCTTCGATTT | atcactggGCCCTGGTGTCTTCGATTT |
| cgcatcaa | TAAGCCAGCACTGCAAGAG | cgcatcaaTAAGCCAGCACTGCAAGAG | GCCCTGGTGTCTTCGATTT | cgcatcaaGCCCTGGTGTCTTCGATTT |
